# Greater Amyloid Burden in Cognitive Networks in Preclinical Alzheimer’s Disease

**DOI:** 10.64898/2026.05.21.726909

**Authors:** Sara A. Nolin, Stephanie Aghamoosa, William J. Rieter, Averi Jones, Paul J Nietert, Andreana Benitez

**Affiliations:** Department of Neurology, Medical University of South Carolina, Charleston, South Carolina, United States; Department of Health Sciences & Research, Medical University of South Carolina, Charleston, South Carolina, United States; Center for Biomedical Imaging, Medical University of South Carolina, Charleston, South Carolina, United States; Department of Radiology and Radiological Sciences, Medical University of South Carolina, Charleston, South Carolina, United States; Department of Public Health Sciences, Medical University of South Carolina, Charleston, South Carolina, United States

**Keywords:** preclinical Alzheimer’s Disease, Alzheimer’s Disease, amyloid-beta, cognition, functional connectivity

## Abstract

**Background:** In preclinical Alzheimer’s disease (pAD), regional patterns of amyloid-β (Aβ) deposition are well characterized but it is unclear how this process varies across functional networks.

**Objective:** Determine how Aβ accumulation in functional networks (“network-amyloid burden” [NAB]) varies by age, network type (cognitive vs. non-cognitive), and Aβ status (Aβ+/Aβ-), and relates to cognition.

**Methods:** 157 cognitively unimpaired adults (45–84 years; n=28 Aβ+ per neuroradiological read) underwent brain MRI, amyloid PET (18F-florbetapir), and neuropsychological testing. NAB was calculated as the mean standard uptake value ratio within 7 networks categorized as cognitive (fronto-parietal, default mode, ventral and dorsal attention, limbic) or non-cognitive (somato-motor, visual). Linear mixed models tested how NAB varies across age, networks (by type and each separately), Aβ status, and their interactions, and relationships between NAB and cognition.

**Results:** NAB increased with age, most prominently in fronto-parietal and default mode networks. NAB was higher in cognitive than non-cognitive networks, and this difference was more pronounced in Aβ+ individuals. NAB was not significantly associated with cognition.

**Conclusions:** Cognitive brain networks are more vulnerable to amyloid accumulation with aging and in pAD than non-cognitive networks. Cognitive NAB may be useful for early detection and as a target for intervention in pAD.

## Introduction

In Alzheimer’s disease (AD), amyloid-β (Aβ) deposition follows a well-established spatial gradient: it begins in higher-order anterior and ventral association cortices and later spreads to primary motor/sensory and subcortical regions^1–3^. Early Aβ accumulation preferentially occurs within metabolically active, highly connected, multimodal brain areas^4^, such as precuneus, medial prefrontal cortex, and posterior cingulate cortex^5,6^. Critically, many of these regions are cortical hubs that are integral to large-scale functional networks that support higher-order cognition^7^. However, because prior work has focused primarily on global and region-level Aβ, the degree to which Aβ deposition varies across functional networks is not well understood. Determining the burden of Aβ in these networks, particularly in the asymptomatic preclinical stage of AD (pAD)^8^, may establish a clear clinico-pathologic correlate of the earliest stages of the disease.

Functional networks can be broadly categorized into cognitive networks (i.e., supporting higher-order cognition) and non-cognitive networks (i.e., primarily involved in somato-sensory functions). In aging and AD, abnormalities in functional connectivity, defined as temporal correlations of activity between brain regions, have been observed within and between cognitive networks, with much research focusing on the default mode network (DMN)^9–11^. It is theorized that the DMN’s high metabolic activity and extensive functional connectivity may render its regions especially vulnerable to early Aβ deposition^12^, contributing to network dysfunction and memory impairment that is characteristic of AD^13,14^.

However, these effects are not isolated to the DMN. Higher global Aβ is linked to connectivity disruptions among regions that overlap with several other cognitive networks, including frontoparietal executive regions, salience-network hubs such as the anterior cingulate, posterior parietal areas of the dorsal attention network, and lateral temporal and occipital association cortices^5,7,12–24^. Additionally, prior work has shown spatial overlap between amyloid deposition and areas belonging to the DMN, frontoparietal, and dorsal and ventral attention networks in pAD^25,26^. To quantify overlap, both studies examined the spatial similarity between functional network maps and amyloid deposition maps, derived from voxel-wise or independent component analysis of PET images. However, neither of these studies measured Aβ burden at the network-level within individuals. This subject-level approach may reveal early network vulnerability in pAD and potentially serve as a marker for early detection and target for intervention.

We therefore developed Network Amyloid Burden (NAB) to quantify and compare Aβ burden in large-scale functional networks. We hypothesized that NAB would: (1) positively correlate with age; (2) be greater in cognitive than non-cognitive networks; (3) more strongly associate with age in cognitive than in non-cognitive networks; (4) exhibit a larger difference between cognitive and non-cognitive networks in neuroradiologically-determined Aβ+ participants than Aβ– participants; and (5) negatively associate with cognitive test performance.

## Methods

### Participants

This study uses baseline data from an ongoing longitudinal observational study of community dwelling, cognitively unimpaired adults. Participants underwent screening to assess eligibility criteria (Table 1) over the phone. In addition to denying self-reported cognitive problems, we excluded participants with normatively low scores on the Montreal Cognitive Assessment (MoCA age and education adjusted *z* ≤−1.0)^27,28^. Lack of objective and self-reported cognitive impairment is consistent with standard criteria for defining cognitively unimpaired individuals in studies of pAD^8,29^. Participants were excluded from analyses for incidental MRI findings (n = 2) or incomplete neuroimaging or neuropsychological data (n = 11). There were 157 participants with complete baseline data (Table 2). Of those, 28 participants (17.8%) were Aβ+ per neuroradiological PET read (described in the *Image acquisition and processing* section below).

**Table 1.** Study Eligibility Criteria.

| Inclusion Criteria | Exclusion Criteria |
| --- | --- |
| Age 45-85 | History of severe/unstable conditions that affect cognition |
| English as first/primary language | Diagnosis of Mild or Major Neurocognitive Disorder |
| Able to provide informed consent | Daily use of medication that adversely impact cognition in aging |
| No self-reported cognitive difficulties | Contraindication to blood draw, PET or MRI |
| Must have co-participant with at least weekly contact | Positive pregnancy test (female participants) |
|  | Non-correctable vision, hearing, or sensorimotor impairments that prevent compliance with study protocol. |
|  | Received radiopharmaceuticals or sufficient ionizing radiation exceeding 50 mSv within previous 12 months. |
|  | Enrolled in a clinical trial or received an investigational medication within the last 30 days |

**Table 2.** Sample characteristics and tests of group differences.

| | Full Sample (N = 157) | | Aβ- Group (n = 129) | | Aβ+ Group (n = 28) | | Group Differences | |
| --- | --- | --- | --- | --- | --- | --- | --- | --- |
|  | Mean (SD) | Range | Mean (SD) | Range | Mean (SD) | Range | stat | p-value |
| **Age (years)** | 67.5 (9.7) | 45.1 – 84.7 | 66.6 (9.5) | 45.06 - 84.7 | 71.6 (9.6) | 47.5 - 84.6 | $U = 1237$ | **.009** |
| **Education (years)** | 15.96 (2.5) | 8 – 25 | 16 (9.5) | 9 – 25 | 16 (9.6) | 8 – 20 | $U = 1769.5$ | .865 |
| **Sex (female; no. and %)** | 108 (68.8%) | - | 89 (69.0%) | - | 19 (67.9%) | - | $\chi^2 = 0$ | 1.0 |
| **Race, ethnicity (no. and %)** | | | | | | | $\chi^2 = 2.4$ | .123 |
| White, Non-Hispanic | 142 (90.4%) | - | 114 (88.4%) | - | 28 (100%) | - | - | - |
| Black/Other, Hispanic/Non-Hispanic | 15 (9.6%) | - | 15 (11.6%) | - | 0 (0%) | - | - | - |
| **General Cognition Composite (Z-score)<sup>a</sup>** | .05 (.4) | -1.3 – 1.01 | .05 (.4) | -1.3 – 1.01 | .04 (.4) | -0.9 – 0.9 | $t(43.5) = 0.12$ | .908 |

<sup>a</sup>Two Aβ - participants were missing complete neuropsychological test data; N = 155, Aβ - n = 127, Aβ + n = 28.

### Image acquisition

Detailed information regarding imaging acquisition has been previously reported in detail^30^. Briefly, participants underwent Aβ PET/CT imaging on a Siemens Biograph mCT Flow PET/CT scanner using ^18^F-florbetapir (Amyvid^TM^) tracer. Each participant was classified as either Aβ+ or Aβ- based on a reading from a nuclear medicine radiologist trained to read Amyvid^TM^ scans. Participants underwent a brain MRI on a research-dedicated 3T Siemens Prisma MRI. The analyses in this paper only used the structural (T1-weighted) image which had the following parameters: acquisition time = 5 minutes and 21 seconds, TR/TE = 2300/2.25ms, voxel size = 1mm^3^, and field of view = 256mm.

### Image processing and analysis

The imaging analysis pipeline is depicted in Figure 1. All image processing and analyses were performed using FSL (FMRIB Software Library, version 6.0.7.13^31^). After preprocessing the T1-weighted image, we registered each participant’s PET image and the 100-parcel, 7-network Schaefer functional network atlas^32,33^ to their T1 (native) space (see Figure 1, Registration). We then created 7 network regions of interest (ROIs) from the native space atlas and masked the network ROIs to only include gray matter. Last, we computed NAB as the mean standard uptake value ratio (mSUVr) within each network, calculated as the mean SUV within each network normalized to the mean SUV of the cerebellum (akin to prior studies computing mSUVr of AD-relevant cortical regions^34^). This resulted in a NAB value for each of the 7 networks. We then computed 1) cognitive NAB by taking the average of NAB values for the cognitive networks (i.e., Fronto-Parietal network [FPN], Default Mode Network [DMN], Ventral Attention Network [VAT], Dorsal Attention Network [DAT], Limbic Network [LIM]) and 2) non-cognitive NAB by taking the average NAB values for Visual (VIS) and Somato-motor (SOM) networks.

**Figure 1.**
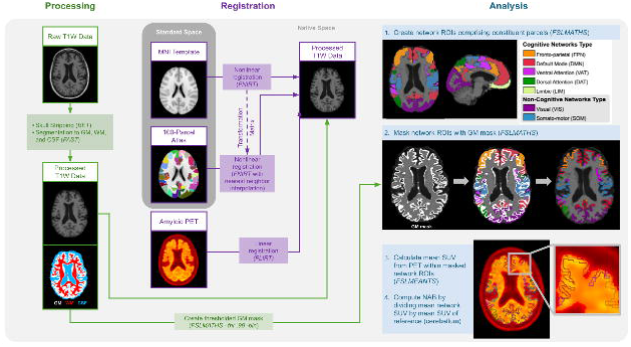

### Neuropsychological testing

Participants were administered the neuropsychological test battery from Version 3 of the National Alzheimer’s Coordinating Centers Uniform Data Set (NACC UDS v3)^35^. Published regression-based norms^35^ were used to compute age, education, and sex-adjusted Z-scores for each test: Craft Story Immediate Recall (paraphrase scoring), Craft Story Delayed Recall (paraphrase scoring), Benson Figure Copy, Benson Figure Recall, Category (Animal) Fluency, Category (Vegetable) Fluency, Multilingual Naming Test, Verbal (F and L) Fluency, Trail Making Test Part A, Trail Making Test Part B, Number Span Forward, and Number Span Backward. Because the present study focuses on amyloid distribution in preclinical AD, where global cognitive performance is often largely preserved, we included a general cognitive composite to index overall functioning while also examining Memory and Executive Functioning domain-specific composites. These domains were selected because prior work has shown they are among the most sensitive to pAD cognitive changes^36^. We computed a general cognition composite score by averaging the z-scores for all tests, the Memory composite score was the average z-score for Immediate Craft Story Recall (paraphrase scoring), Delayed Craft Story Recall (paraphrase scoring), and the Benson Figure Recall, the Executive Functioning composite score was the average z-score for Trail Making Test Part B and Number Span Backward^37^.

### Statistical Analysis

All statistical analyses were performed using SAS® v9.4 (SAS Institute, Cary, NC). We used linear mixed models (LMMs) to test for differences in NAB across age, networks (cognitive vs. non-cognitive and the 7 networks separately), Aβ group, and their interactions. All models were estimated using Restricted Maximum Likelihood (REML) and included a fixed effect for age as a continuous covariate and random intercepts for participants. Repeated measures were modeled using an unstructured covariance matrix, which demonstrated superior model fit relative to models with compound symmetry covariance structures.

Effect sizes^38^ were calculated by standardizing model-estimated effects using a common between-person standard deviation (SD= 0.0735), which was estimated based on an LMM that decomposed total variance into between- and within-person components. The numerators for each of the effect sizes were based on model-estimated contrasts. Models included one or more of the following categorical fixed effects: “network type” had two levels (cognitive, non-cognitive), “7 networks” had seven levels (FPN, DMN, VAT, DAT, LIM, SOM, VIS where VIS was set as the reference since it is a sensory network that is anatomically far from areas with early amyloid accumulation), “Aβ group” had two levels (Aβ+, Aβ-). We fit 5 different models (with NAB as the dependent variable in each model) to investigate our hypotheses. Model 1 included main effects for age and “network type”.

Model 2 included age, “network type”, and their interaction. Model 3 included age, “7 networks”, and their interaction. Model 4 included age, Aβ group, “network type”, and their interaction. Model 5 included age, Aβ group, “7 networks”, and their interaction. For each LMM, a likelihood ratio test compared goodness of fit of the full model vs. a null model containing only the intercept. All models had a significantly better fit from the null model. For LMMs including all 7 networks, pairwise comparisons with Tukey HSD adjustment tested for differences between each of the networks. We evaluated the association between cognitive NAB and cognition (general cognition, Memory, and Executive Functioning composite z-scores) using partial Spearman’s rank correlation, controlling for age.

### Ethics

The study was approved by the Medical University of South Carolina institutional review board. All participants provided written informed consent.

## Results

### Differences in NAB by Age and Network Type

NAB was positively associated with age (B= .003, SE= .001, *p* = .012, *d* = 0.04), such that for every year increase in age, NAB increased by 0.0026 (Table S1). NAB was higher in cognitive networks than in non-cognitive networks (B= .055, SE= .07, *p* < .001, *d* = 0.74; Table S1, Figure 2). However, the interaction with age was not significant, indicating that the magnitude of the difference in NAB between cognitive and non-cognitive networks did not vary significantly with age (*p* = .203; Table S2). NAB in cognitive networks was not significantly associated with general cognition (*r*_s_ = −0.071, *p* = .381), memory (*r*_s_ = −0.02, *p* = .776), or executive functioning(*r*_s_ = −0.02, *p* = .791).

**Figure 2.**
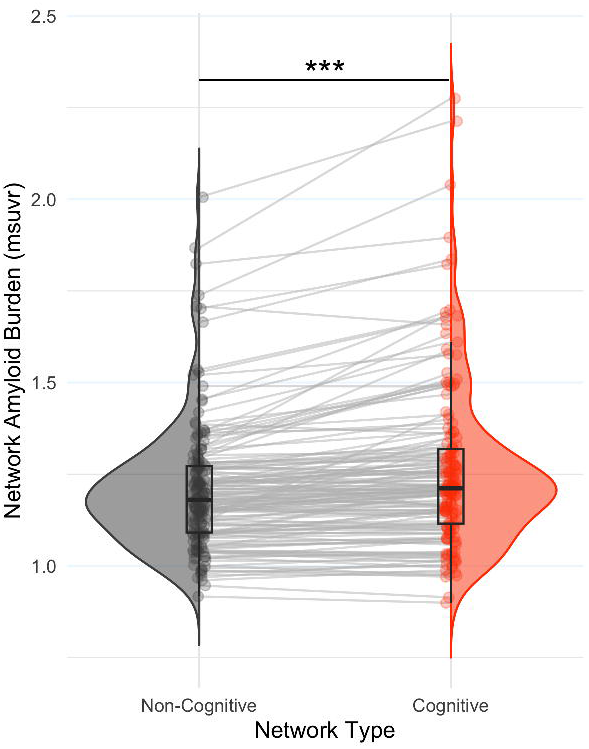

### Differences in NAB across the 7 Networks

NAB differed significantly across the 7 networks when controlling for age, *F*(6, 155) = 3.70, *p* = .0018 (Table S3). Model-estimated mean NAB within each network at the mean age of the sample (i.e., age-adjusted least squares means) are presented in Figure 3A. Post hoc Tukey-Kramer comparisons between networks indicated that compared to SOM, all networks had significantly higher NAB. Additionally, NAB was significantly higher in DMN (Δ = 0.01818, *p_adj_* = 0.0171), FPN (Δ = 0.02092, *p_adj_* = 0.0060), and VAT (Δ = 0.02381, *p_adj_* < 0.0001) than DAT. There was also a significant age X “7 network” interaction, *F*(6, 155) = 3.91, *p* = .0012, indicating that the association between age and NAB differed across the seven networks. This effect was strongest for FPN and DMN, such that these two networks showed a significantly greater increase in NAB with age compared to the reference network, VIS (Figure 3, Table S3).

**Figure 3.**
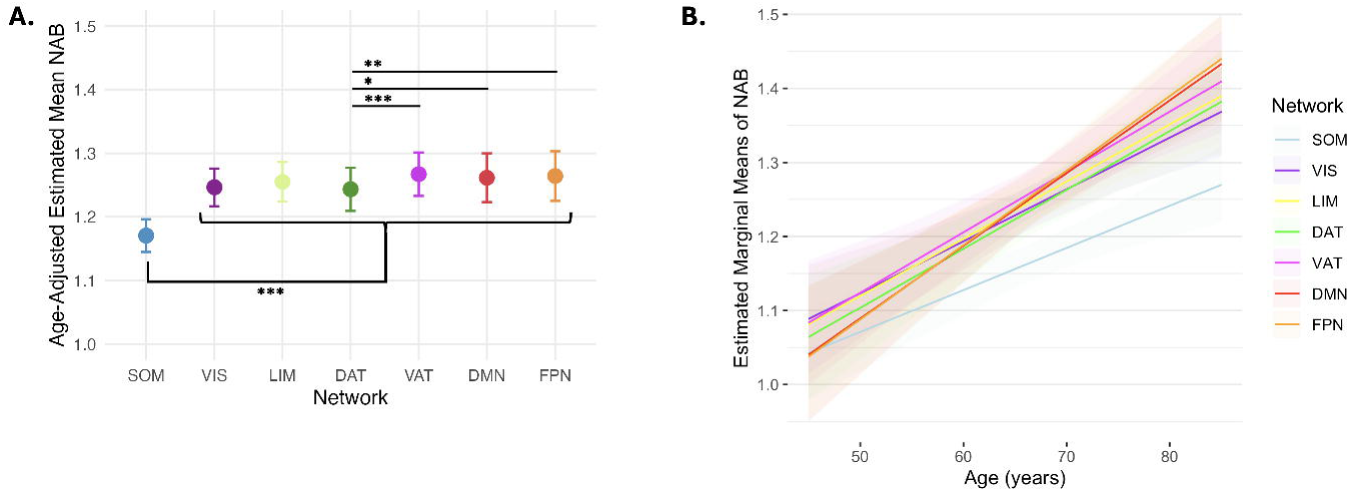

### Differences in NAB by Aβ Status and across Networks

NAB was higher overall in the *Aβ*+ than *Aβ* – group, *F*(1,154) = 56.84, *p* < .0001. There was a group X network type interaction, whereby the effect of higher NAB in cognitive vs. non-cognitive networks was greater in the *Aβ*+ than *Aβ*- group (Figure 4A and Table S4). There was also a significant group X “7 networks” interaction, *F*(6, df) = 29.53, *p* < .0001, such that the effect of higher NAB in the *Aβ*+ vs. *Aβ*- group was strongest in the following networks (in order of decreasing effect size): FPN, DMN, DAT, VAT (Figure 4B and Table S5).

**Figure 4.**
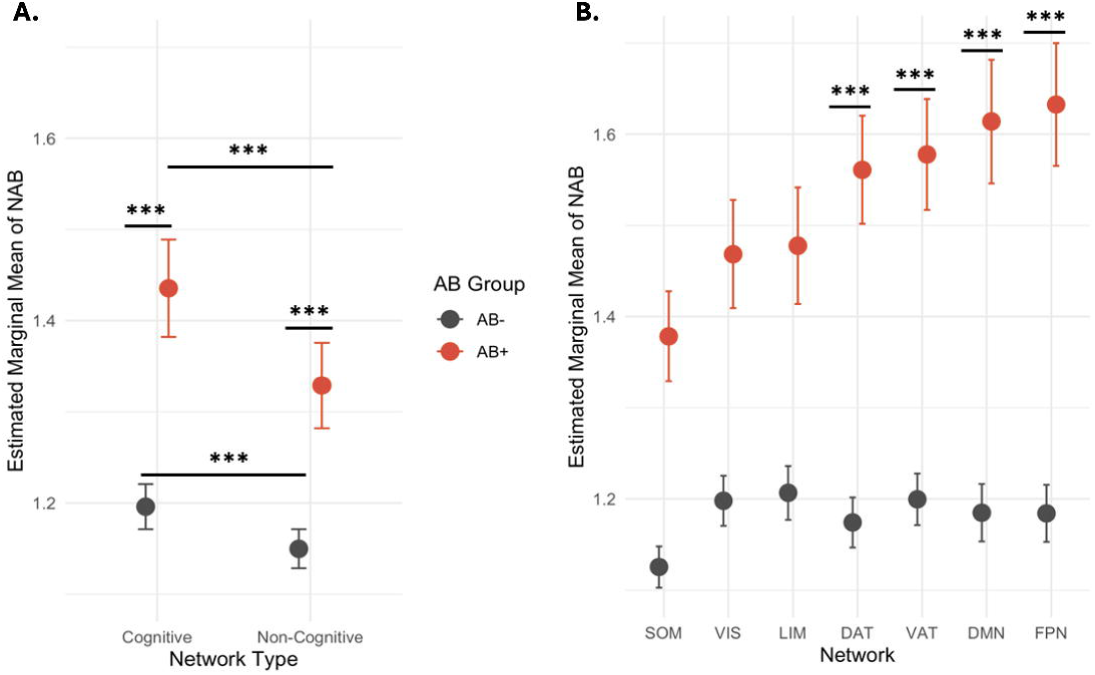

## Discussion

This study investigated how amyloid deposition in functional brain networks vary by age and Aβ status in cognitively unimpaired adults. Using a novel metric of network amyloid burden (NAB), we showed that NAB correlated with age, was higher in cognitive than non-cognitive networks, and exhibited the strongest age-related associations in select cognitive networks. Additionally, the effect of higher NAB in cognitive vs. non-cognitive networks was greater in Aβ+ than Aβ-individuals, suggesting greater network-level vulnerability in pAD. However, cognitive NAB was not associated with cognitive composite scores in this cognitively unimpaired sample.

This study extends prior work by quantifying amyloid burden within large-scale functional networks, revealing differential amyloid accumulation across networks. Whereas prior work has evaluated the spatial overlap of amyloid deposition with functional networks^6,25,26^, our approach allowed us to directly investigate the degree to which amyloid burden varies across networks. We found significantly higher NAB overall in cognitive networks compared to non-cognitive networks. Across the seven canonical networks, NAB was elevated in all networks when compared to the somato-motor network, with the strongest effects in the default mode, fronto-parietal, and ventral attention networks.

Additionally, the default mode and frontoparietal networks showed the greatest age-related increases in NAB compared to the visual network. The elevated NAB observed in these networks aligns with prior work demonstrating that Aβ preferentially accumulates in highly connected, metabolically active regions, particularly within the default mode network^39–41^. Together, these findings support a network-based framework for quantifying early Aβ deposition in pAD.

Critically, we also found that this differential pattern of Aβ deposition across networks varied according to neuroradiologically-defined Aβ status. Specifically, the effect of higher amyloid burden in cognitive than non-cognitive networks was more pronounced in the Aβ+ group (i.e., those with pAD) than the Aβ- group, and this group difference was most pronounced in frontoparietal, default mode, and dorsal and ventral attention networks. That is, beyond an expected global elevation in amyloid in Aβ+ individuals, there was a larger difference between cognitive and non-cognitive NAB in Aβ+ compared to Aβ-individuals. This selective network vulnerability in pAD suggests that cognitive networks may serve as potential targets for early intervention and disease monitoring. These findings support the idea that cognitive NAB may be a sensitive biomarker of the AD pathological process. Importantly, the NAB method extends beyond prior spatial overlap approaches, providing a straightforward, quantitative measure of network-level Aβ burden on an individual level.

The absence of an association between cognitive NAB and cognitive test performance likely reflects multiple factors. It is possible that Aβ burden quantified at the network-level may not be sensitive to early, subtle cognitive deficits in the pAD stage. Alternatively, the cognitive measures used in this study (i.e., composites derived from the NACC neuropsychological battery), may not have been sensitive to subtle cognitive deficits that are present in the pAD stage. Although composite measures demonstrate better sensitivity to cognitive deficits and AD biomarkers in pAD than single test scores^36,42^, there is no consensus regarding a gold-standard measure for assessing cognition in pAD. In this study, scores on the general cognitive composite did not differ by amyloid status and were limited in range to roughly 1 SD below and above the normative mean (see Table 2), suggesting either limited sensitivity of the measure or no true differences in cognition between groups. It is possible that individuals in the Aβ+ group were still early in the disease process and that amyloid burden had not risen sufficiently for detectable cognitive symptoms to present^43^. Future work could include longitudinal data to evaluate change in NAB over time and investigate whether it predicts subtle cognitive decline using measures sensitive to the preclinical stage.

There are several limitations of this study that point to important future directions. First, the sample is relatively socio-demographically homogenous and Aβ+ constituted only 17% of total sample, so replication of our findings in larger and more representative samples is needed to ensure generalizability to the broader population. Second, our results could depend on the analytic choices that influence how the PET signal is assigned to a network prior to calculating NAB (e.g., atlas selection, registration, spatial smoothing, gray-matter masking, etc.). It will be critical for future work to transparently report analytic methods to facilitate replication and extension of these findings. A key future direction is to examine how NAB relates to functional network connectivity measured with fMRI, at rest or during cognitive tasks. Prior work shows that connectivity alterations, especially in cognitive networks, precede cognitive impairment^44^ and can predict subsequent cognitive decline^45^, suggesting that combining network-level amyloid measures with connectivity analyses could clarify early disease mechanisms and improve predicative utility. Although tau accumulation is minimal in preclinical stages, it would be informative to also assess whether similar network-specific patterns of tau emerge in later stages of AD.

Taken together, these findings reveal early vulnerability of large-scale cognitive networks to amyloid accumulation in pAD. NAB may provide a promising approach for quantifying the network-level expression of Aβ, with the potential to contribute to early detection, mechanistic understanding, and intervention targeting in future work.

## Acknowledgments

We thank K. Madden, K. Thorn, S. Helton, and J. Kellam for their help with participant recruitment, data collection, and data entry in support of this project. We offer our utmost appreciation to all our participants, whose enthusiasm for research is unrivaled and deeply valued by our team.

## Author Contributions

Sara A Nolin (Conceptualization, Data curation, Investigation, Methodology, Formal Analysis, Writing – Original Draft, Visualization); Stephanie Aghamoosa (Data curation, Methodology, Formal Analysis, Visualization, Writing- Original Draft); William J. Rieter (Investigation, Writing- Review & Editing); Averi Jones (Data curation, Methodology, Writing-Review & Editing); Paul J Nietert (Formal Analysis, Writing- Review & Editing); Andreana Benitez (Funding Acquisition, Project Administration, Investigation, Supervision, Writing-Review & Editing).

## Statements and Declarations

### Ethical considerations

The study was approved by the Medical University of South Carolina institutional review board (#73604) on 12/19/2017.

### Consent to participate

All participants provided informed written consent.

### Consent for publication

Not applicable.

### Data availability

Study data can be shared upon reasonable request, contingent upon compliance with applicable institutional policies, regulatory requirements, and any relevant data use agreements. Code used for processing and statistical analysis is available via: https://github.com/muscbridge/Amyloid_Network/.

