## Supplementary material for "Greater Amyloid Burden in Cognitive Networks in Preclinical Alzheimer’s Disease": Table S1 (from another version of this preprint: version 1, 2026-05-26)

### Supplemental Tables

**Table S1**

*Linear Mixed Model Results for Age and Network-Type Predictors of Network Amyloid Burden (N=157).*

| Predictors | Estimate | SE | t | 95% CI (Lower) | 95% CI (Upper) | p | Effect Size |
|---|---|---|---|---|---|---|---|
| Intercept | 1.0005 | 0.0698 | 14.3400 | 0.8627 | 1.1383 | <.0001 | - |
| **Network Type (Cognitive)** | **0.0547** | **0.0037** | **14.8700** | **0.0475** | **0.0620** | **<.0001** | **0.74** |
| **Age** | **0.0026** | **0.0010** | **2.5300** | **0.0006** | **0.0046** | **0.0123** | **0.04** |

*Note*. *SE* = Standard Error; *CI* = Confidence Interval. Effect sizes were calculated by standardizing model-estimated effects using a common between-person standard deviation (SD = 0.0735), estimated from a general linear model decomposing total variance into between- and within-person components. For models without interactions, effect sizes correspond to standardized fixed-effect estimates; for models including interactions, effect sizes correspond to standardized model-estimated contrasts.

**Table S2**

*Linear Mixed Model Results for Age-by-Network-type Predictors of Network Amyloid Burden (N=157).*

| Predictor | Estimate | SE | t | 95% CI (Lower) | 95% CI (Upper) | p |
|---|---|---|---|---|---|---|
| Intercept | 0.9649 | 0.0753 | 12.8200 | 0.8162 | 1.1135 | <.0001 |
| Network Type (Cognitive) | 0.0222 | 0.0261 | 0.8500 | -0.0293 | 0.0737 | 0.3954 |
| **Age** | **0.0031** | **0.0011** | **2.8300** | **0.0009** | **0.0053** | **0.0053** |
| Age * Network Type (Cognitive) | 0.0005 | 0.0004 | 1.2800 | -0.0003 | 0.0012 | 0.2025 |

*Note*. *SE* = Standard Error; *CI* = Confidence Interval.

**Table S3**

*Linear Mixed Model Results for Age-by-Network Predictors of Network Amyloid Burden (N=157).*

| Predictor | Estimate | SE | t | 95% CI (Lower) | 95% CI (Upper) | p | Effect Size |
|---|---|---|---|---|---|---|---|
| Intercept | 0.7729 | 0.1039 | 7.4400 | 0.5676 | 0.9781 | <.0001 | - |
| **Age** | **0.0070** | **0.0015** | **4.6000** | **0.0040** | **0.0100** | **<.0001** | **-** |
| Network | | | | | | | |
| Visual | - | | | | | | |
| Somato-motor | 0.0137 | 0.0509 | 0.2700 | -0.0869 | 0.1143 | 0.7887 | - |
| Limbic | -0.0396 | 0.0534 | -0.7400 | -0.1451 | 0.0659 | 0.4593 | - |
| Ventral Attn. | -0.0567 | 0.0649 | -0.8700 | -0.1849 | 0.0715 | 0.3840 | - |
| Dorsal Attn. | -0.0672 | 0.0541 | -1.2400 | -0.1740 | 0.0396 | 0.2159 | - |
| **Fronto-parietal** | **-0.1888** | **0.0724** | **-2.6100** | **-0.3318** | **-0.0458** | **0.0100** | **-** |
| **Default Mode** | **-0.1752** | **0.0656** | **-2.6700** | **-0.3048** | **-0.0456** | **0.0084** | **-** |
| Age * Network | | | | | | | |
| Visual | - | | | | | | |
| Somato-motor | -0.0013 | 0.0007 | -1.7700 | -0.0028 | 0.0002 | 0.0780 | 0.08 |

| | | | | | | | |
|---|---|---|---|---|---|---|---|
| Limbic | 0.0007 | 0.0008 | 0.9200 | -0.0008 | 0.0023 | 0.3614 | 0.11 |
| Ventral Attn. | 0.0011 | 0.0010 | 1.2100 | -0.0007 | 0.0030 | 0.2300 | 0.11 |
| Dorsal Attn. | 0.0010 | 0.0008 | 1.2000 | -0.0006 | 0.0025 | 0.2326 | 0.11 |
| **Fronto-parietal** | **0.0031** | **0.0011** | **2.8800** | **0.0010** | **0.0052** | **0.0045** | **0.14** |
| **Default Mode** | **0.0028** | **0.0010** | **2.9300** | **0.0009** | **0.0047** | **0.0039** | **0.13** |

*Note. SE* = Standard Error; *CI* = Confidence Interval. Effect sizes were calculated by standardizing model-estimated effects using a common between-person standard deviation (SD = 0.0735), estimated from a general linear model decomposing total variance into between- and within-person components. For models without interactions, effect sizes correspond to standardized fixed-effect estimates; for models including interactions, effect sizes correspond to standardized model-estimated contrasts.

**Table S4**

*Linear Mixed Model Results for Age, PET Read, and Network-type Predictors of Network Amyloid Burden (N=157).*

| Predictor | Estimate | SE | t | 95% CI (Lower) | 95% CI (Upper) | p | Effect Size |
|---|---|---|---|---|---|---|---|
| Intercept | 0.9705 | 0.0677 | 14.3300 | 0.8367 | 1.1043 | <.0001 | - |
| **Network Type (Cognitive)** | **0.0462** | **0.0038** | **12.2000** | **0.0387** | **0.0536** | **<.0001** | **-** |
| **Age** | **0.0027** | **0.0010** | **2.6500** | **0.0007** | **0.0046** | **0.0089** | **-** |
| **PET Read (Aβ+)** | **0.1790** | **0.0262** | **6.8300** | **0.1273** | **0.2308** | **<.0001** | **-** |
| **Aβ+ * Cognitive** | **0.0605** | **0.0090** | **6.7600** | **0.0428** | **0.0782** | **<.0001** | **3.26** |

Note. SE = Standard Error; CI = Confidence Interval. Effect sizes were calculated by standardizing model-estimated effects using a common between-person standard deviation (SD = 0.0735), estimated from a general linear model decomposing total variance into between- and within-person components. For models without interactions, effect sizes correspond to standardized fixed-effect estimates; for models including interactions, effect sizes correspond to standardized model-estimated contrasts.

**Table S5**

*Linear Mixed Model Results for PET-Read-by-Network Predictors of Network Amyloid Burden (N=157).*

| Predictor | Estimate | SE | t | 95% CI (Lower) | 95% CI (Upper) | p | Effect Size |
|---|---|---|---|---|---|---|---|
| Intercept | 1.0186 | 0.0683 | 14.9200 | 0.8837 | 1.1535 | <.0001 | - |
| **Age** | **0.0027** | **0.0010** | **2.6500** | **0.0007** | **0.0046** | **0.0089** | **-** |
| **PET Read (Aβ+)** | **0.2705** | **0.0332** | **8.1600** | **0.2050** | **0.3360** | **<.0001** | **-** |
| Network | | | | | | | |
| Visual | - | | | | | | - |
| **Somato-motor** | **-0.0727** | **0.0080** | **-9.1000** | **-0.0884** | **-0.0569** | **<.0001** | **-** |
| Limbic | 0.0087 | 0.0083 | 1.0400 | -0.0078 | 0.0251 | 0.2996 | - |
| Ventral Attn. | 0.0015 | 0.0095 | 0.1600 | -0.0172 | 0.0202 | 0.8711 | - |
| Dorsal Attn. | -0.0238 | 0.0075 | -3.1800 | -0.0385 | -0.0090 | 0.0018 | - |
| Fronto-parietal | -0.0139 | 0.0099 | -1.4100 | -0.0333 | 0.0056 | 0.1619 | - |
| Default Mode | -0.0132 | 0.0090 | -1.4600 | -0.0309 | 0.0046 | 0.1461 | - |
| AB+ * Network | | | | | | | |

| | | | | | | | |
|---|---|---|---|---|---|---|---|
| Visual | - | | | | | | - |
| Somato-motor | -0.0174 | 0.0189 | -0.9200 | -0.0548 | 0.0199 | 0.3575 | 3.44 |
| Limbic | 0.0006 | 0.0197 | 0.0300 | -0.0384 | 0.0396 | 0.9762 | 3.69 |
| **Ventral Attn.** | **0.1078** | **0.0224** | **4.8100** | **0.0636** | **0.1521** | **<.0001** | **5.15** |
| **Dorsal Attn.** | **0.1162** | **0.0177** | **6.5600** | **0.0812** | **0.1512** | **<.0001** | **5.26** |
| **Fronto-parietal** | **0.1780** | **0.0234** | **7.6200** | **0.1318** | **0.2241** | **<.0001** | **6.10** |
| **Default Mode** | **0.1586** | **0.0213** | **7.4400** | **0.1165** | **0.2007** | **<.0001** | **5.84** |

*Note.* *SE* = Standard Error; *CI* = Confidence Interval. Effect sizes were calculated by standardizing model-estimated effects using a common between-person standard deviation (SD = 0.0735), estimated from a general linear model decomposing total variance into between- and within-person components. For models without interactions, effect sizes correspond to standardized fixed-effect estimates; for models including interactions, effect sizes correspond to standardized model-estimated contrasts.
